# SPiQE: an automated analytical tool for detecting and characterising fasciculations in amyotrophic lateral sclerosis

**DOI:** 10.1101/571893

**Authors:** J Bashford, A Wickham, R Iniesta, E Drakakis, M Boutelle, K Mills, C Shaw

**Affiliations:** Department of Basic and Clinical Neurosciences, King’s College London; Department of Bioengineering, Imperial College London; Department of Biostatistics and Health Informatics, King’s College London

**Keywords:** Amyotrophic lateral sclerosis, fasciculation, high-density surface EMG, biomarker

## Abstract

**OBJECTIVES:** Fasciculations are a clinical hallmark of amyotrophic lateral sclerosis (ALS). Compared to concentric needle EMG, high-density surface EMG (HDSEMG) is non-invasive and records fasciculation potentials (FPs) from greater muscle volumes over longer durations. To detect and characterise FPs from vast data sets generated by serial HDSEMG, we developed an automated analytical tool.

**METHODS:** Six ALS patients and two control patients (one with benign fasciculation syndrome and one with multifocal motor neuropathy) underwent 30-minute HDSEMG from biceps and gastrocnemius monthly. In MATLAB we developed a novel, innovative method to identify FPs amidst fluctuating noise levels. One hundred repeats of 5-fold cross validation estimated the model’s predictive ability.

**RESULTS:** By applying this method, we identified 5,318 FPs from 80 minutes of recordings with a sensitivity of 83.6% (+/-0.2 SEM), specificity of 91.6% (+/-0.1 SEM) and classification accuracy of 87.9% (+/-0.1 SEM). An amplitude exclusion threshold (100μV) removed excessively noisy data without compromising sensitivity. The resulting automated FP counts were not significantly different to the manual counts (p=0.394).

**CONCLUSION:** We have devised and internally validated an automated method to accurately identify FPs from HDSEMG, a technique we have named Surface Potential Quantification Engine (SPiQE).

**SIGNIFICANCE:** Longitudinal quantification of fasciculations in ALS could provide unique insight into motor neuron health.

**Highlights:**

- SPiQE combines serial high-density surface EMG with an innovative signal-processing methodology
- SPiQE identifies fasciculations in ALS patients with high sensitivity and specificity
- The optimal noise-responsive model achieves an average classification accuracy of 88%

## 1 Introduction

ALS is caused by the progressive dysfunction and death of motor neurons and affects ~1,200 people in the UK every year.(Al-Chalabi and Hardiman, 2013) It typically causes progressive paralysis and death within three to five years of symptom onset. There is only one licensed drug in Europe (riluzole) with modest survival benefit.(Bensimon et al., 1994) Drug trials in ALS are time-consuming for patients, expensive for funders and hampered by insensitive measurements of disease progression. Consequently, there is a drive to discover novel biomarkers that could be incorporated into clinical trials to expedite drug discovery.(Benatar et al., 2016)

A motor unit (MU) comprises the motor neuron cell body, axon, terminal branches and connecting muscle fibres. Ailing motor neurons are electrically unstable and spontaneously discharge electrical impulses causing fasciculation potentials (FPs).(de Carvalho and Swash, 2016b) Subsequently, the motor neuron becomes electrically unresponsive and dies, disrupting MU architecture through denervation. Orphaned muscle fibres can become re-innervated by sprouting motor axons. This process of denervation and reinnervation results in MU potentials becoming longer in duration with more complex morphologies over time.(Conradi et al., 1982, de Carvalho and Swash, 2013)

These neurophysiological changes can be observed as a brief snapshot by concentric NEMG, which involves the insertion of a fine needle into muscles to record FPs.(Mills, 2010) This detects electrical activity within a small field comprising approximately 200 muscle fibres and can be painful, making repeated or extended recordings undesirable. Furthermore, there is little chance of recording from the same MU serially. An alternative approach is to use HDSEMG, where a grid of non-invasive sensors is applied to the skin.(Mateen et al., 2008, Howard and Murray, 1992) FPs can be recorded for longer, covering a greater volume of muscle, and can be repeated at regular intervals.(de Carvalho and Swash, 2016a)

One of the biggest challenges facing surface EMG analysis is noise.(De Luca et al., 2010) This arises from both biological and technical factors, including the effects of soft tissue, poor skin-electrode contact and contamination from external electrical sources. This is likely to vary considerably between patients, anatomical sites and recording visits. Previous authors have used a constant threshold for FP detection ranging from 50-100μV.(de Carvalho and Swash, 2013, de Carvalho and Swash, 2016a) We set out to explore the optimal analytical pipeline to account for varying noise levels.

The unique properties of serial HDSEMG predict an ability to characterise the number, temporal patterns and morphology of FPs over time. In order to capitalise on this, it was necessary to devise an automated system capable of processing large data sets in a systematic and robust way. This system, which we have named Surface Potential Quantification Engine (SPiQE), is accurately able to identify FPs from raw HDSEMG recordings as a measure of motor neuron health and may act as a biomarker of disease progression.

## 2 Methodology

### 2.1 Patient characteristics

Six patients with ALS and two control patients (one with benign fasciculation syndrome and one with multifocal motor neuropathy) underwent 42 assessments in total at intervals of at least one month (*table 1*). ALS patients were diagnosed with probable or definite ALS using the revised El Escorial Criteria.(Brooks et al., 2000) Ethical approval was obtained from the North of Scotland Research Ethics Service (Ref: 15/NS/0103). Patients were recruited from the King’s College Hospital Motor Nerve Clinic between Jan-Feb 2016.

**Table 1.** Patient characteristics. ALS, amyotrophic lateral sclerosis; MMN, multifocal motor neuropathy; BFS, benign fasciculation syndrome. Muscle powers were assessed at baseline by the same clinician (JB).

| Patient No. | Age (years) | Gender | Diagnosis | Site of symptom onset | Duration since symptom onset (months) | Biceps power (MRC scale, 5=normal) |  | Gastrocnemius power (MRC scale, 5=normal) |  | Number of assessments undertaken |
| --- | --- | --- | --- | --- | --- | --- | --- | --- | --- | --- |
|  |  |  |  |  |  | R | L | R | L |  |
| 1 | 59 | M | ALS | Left leg | 24 | 5 | 5 | 5 | 5 | 4 |
| 2 | 57 | M | ALS | Right arm | 19 | 4 | 4 | 4 | 4 | 5 |
| 3 | 50 | M | ALS | Left arm | 23 | 2 | 2 | 5 | 5 | 4 |
| 4 | 58 | M | ALS | Right arm | 60 | 1 | 1 | 5 | 5 | 6 |
| 5 | 59 | F | ALS | Bulbar | 27 | 5 | 5 | 5 | 5 | 6 |
| 6 | 61 | M | ALS | Left arm | 10 | 5 | 5 | 5 | 5 | 7 |
| 7 | 68 | F | MMN | Left arm | 204 | 4 | 5 | 5 | 5 | 6 |
| 8 | 57 | M | BFS | Both legs | 8 | 5 | 5 | 5 | 5 | 4 |

### 2.2 Data collection

At each assessment, 30-minute HDSEMG recordings were taken from biceps brachii and medial gastrocnemii bilaterally (left-sided muscles were recorded simultaneously followed by right-sided muscles). The sensor had 64 circular electrodes (8×8 grid; electrode diameter 4.5mm; inter-electrode distance 8.5mm; TMS International BV, The Netherlands). Linear measurements between the medial inferior corner of the grid and the medial epicondyle (biceps) or malleolus (gastrocnemius) guided sensor placement on subsequent visits. The skin was lightly scrubbed with an abrasive gel and a 70% alcohol wipe. A template facilitated the application of conducting gel. Reference electrodes (3×5cm) were placed over the ipsilateral olecranon (biceps) and dorsum of the foot (gastrocnemius). Patients relaxed on the examination couch with legs in a horizontal, partially flexed position and forearms prone with an elbow angle of 90-120 degrees.

The signals were amplified by the Refa-64 EMG Recording System (TMS International BV, The Netherlands). The difference between the average signal from all 64 channels and the reference electrode was subtracted from each channel. The raw HDSEMG data was stored as a proprietary Polybench file at a sampling rate of 2048Hz per channel. A 30-minute recording used approximately 1.1GB of computer storage space. Sensors were cleaned using propran-2-ol solution in the laboratory and re-used up to three times according to manufacturer guidance.

### 2.3 Computation and statistical analysis

All FP computation was performed in MATLAB (R2014a) using specifically designed scripts. Statistical tests were performed in Prism V7.0a. The Mann-Whitney and Wilcoxon signed-rank tests were used for non-parametric data. Laptops with Intel i7 (2.5GHz) processor were used for all analysis.

### 2.4 Data processing

Each 30-minute recording comprised 64 channels. The amplitude was recorded at intervals of ~0.5ms. A bandpass filter (20-500Hz) was applied without notch filtering. The perimeter channels were excluded as these had the poorest skin contact. Amongst the remaining 36 channels, poorly behaving channels were excluded in an automated way. This comprised channels that were null due to absent electrical contact or those that contained excessive noise, artefacts or baseline drift. A detailed description of these methods is beyond the scope of this manuscript, but they involved performing the fast fourier transform, analysing the area under the curve and calculating the amplitude range across the channel. Channels that fell outside the 95% confidence interval for any of these parameters were excluded.

We have developed a novel spike detection mechanism based on a probabilistic analysis of spikes in relation to baseline noise. Briefly, spikes were detected based on a probability threshold of >98% (*figure 1a*). A single super-channel was created by merging spikes from different channels based on the maximum amplitude for each spike (*figure 1b).* Spatial information was retained by recording the channel of origin for each spike. A quality assurance process ensured each recorded spike represented a single spike. A train of at least four FPs with inter-FP intervals (IFIs) <250ms was designated as voluntary activity.

**Figure 1.**
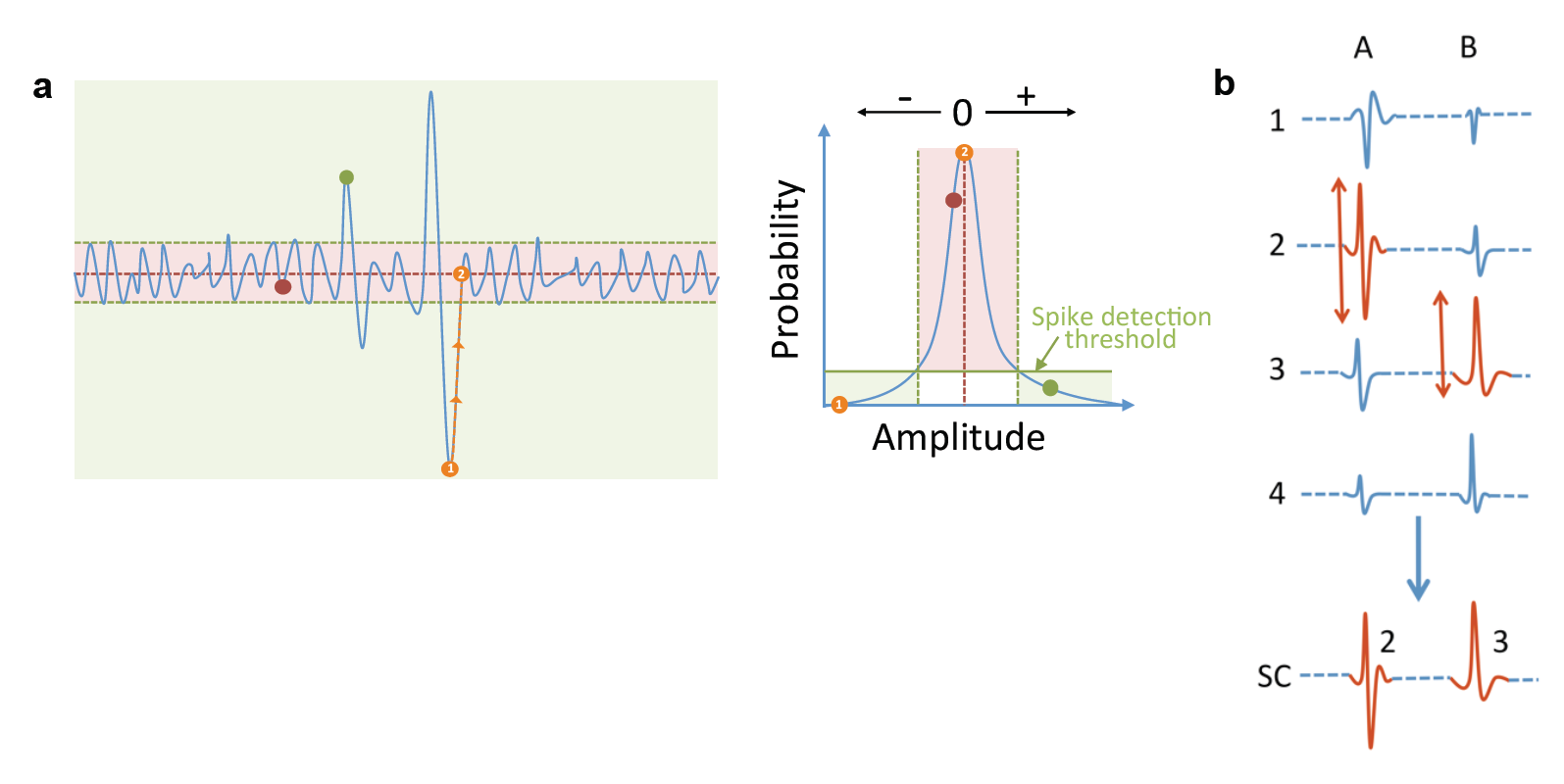
Data processing. **a** *Probability threshold.* The spike detection threshold was based on the probability of a given amplitude occurring in the thirty-minute recording. A spike’s duration was found by tracking the spike to its nearest zero point (*orange circles*). Adjacent phases (positive-negative transition) were joined and recorded as a single spike. **b** *Principle of the super-channel (SC).* For spikes A and B, the channel (1-4) with the highest peak-trough amplitude (shown in red) was transferred into the SC. The channel of origin for each spike was stored.

### 2.5 Optimisation and validation of the analytical pipeline

The aim of the automated pipeline was to identify which spikes were true FPs. The main challenge in this setting concerned fluctuating noise levels during and between recordings. If not appropriately managed, this could introduce significant inaccuracies in FP counts. Noisy data could introduce errors in two main ways. First, if the amplitude threshold for FP inclusion was set too low relative to the noise level, erroneous FPs would be counted, significantly reducing the pipeline’s specificity. Second, if there were sustained periods of excessively noisy and poorly interpretable data that were not excluded, then the time denominator would be inappropriately high, thereby underestimating FP frequencies.

For these reasons, two amplitude thresholds were necessary for FPs. The first, coined the amplitude inclusion threshold (AT_inc_), signified the amplitude above which FPs would be included (phases 1-3). The second, coined the amplitude exclusion threshold (AT_exc_), acted as a trigger for exclusion of excessively noisy data from the analysis (phase 4).

#### 2.5.1 Phase one - Relationships between manual FP counts, noise levels and optimal amplitude inclusion thresholds

Visits 1-4 were used for validation to ensure equal contributions from each patient. For each of the 32 assessments, five one-minute sample recordings starting at 5/10/15/20/25 minutes underwent manual FP counts. The raw data were viewed in consecutive ten-second windows. All manual counts were performed by one assessor (JB) before automated processing. Pre-defined rules for FP inclusion were used:

a. The autoscale function in Polybench optimised FP visualization;
b. An FP was defined as a spike in ≥10 channels (/64) simultaneously;
c. An FP was excluded if part of a train of similar spikes representing voluntary activity.

Automated analysis was performed on the same one-minute recordings. The AT_inc_ was varied to find the lowest value that produced an automated FP count equal to the manual count (+/-1), which was labeled the optimal AT_inc_.

Average noise bands were calculated for each detected spike (*figure 2a*). This was defined as the difference between the mean positive amplitude and the mean negative amplitude for one second either side of the spike. Mean noise bands were calculated for each of the one-minute recordings and plotted against the corresponding optimal AT_inc_. A best-fit line was calculated by the weighted least squares regression method due to heteroscedasticity of the data (*figure 2b*).

**Figure 2.**
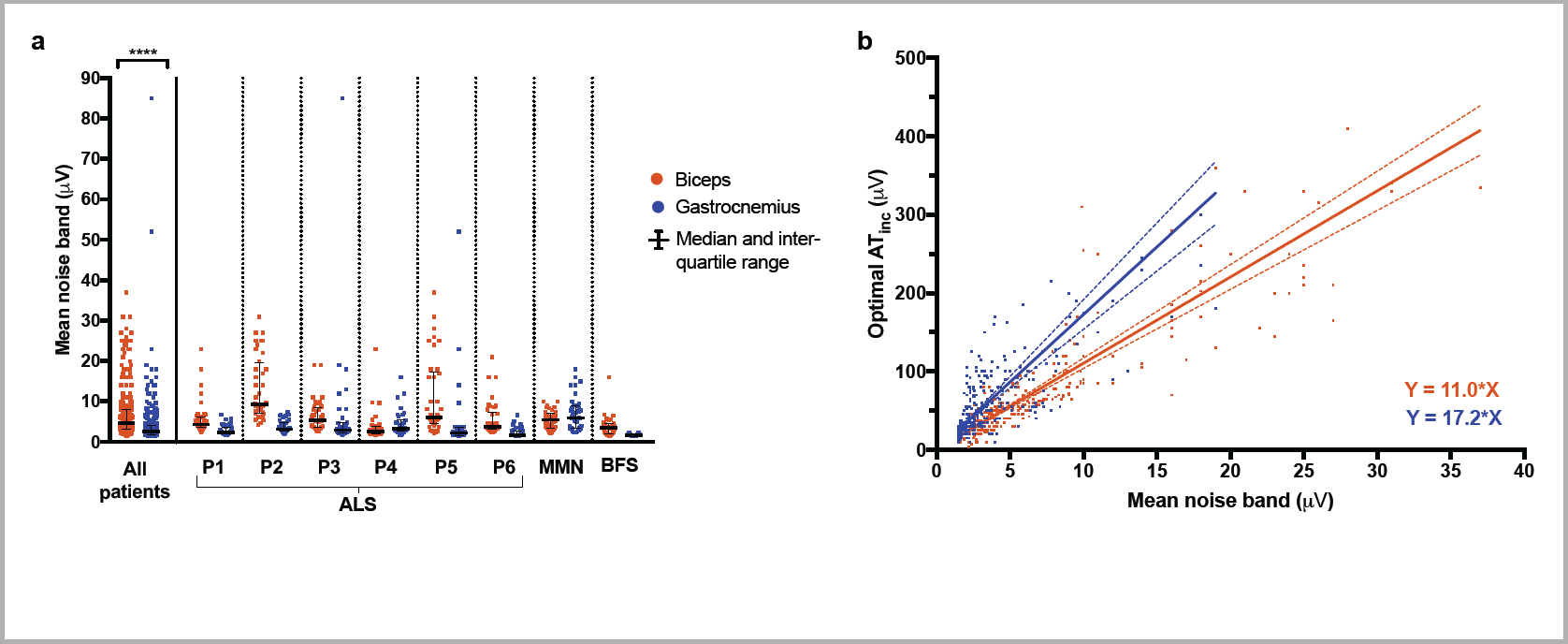
Noise level analysis (phase one). **a** *Distribution of mean noise bands for biceps and gastrocnemius.* Each data point represents one minute of recording. There was a significant difference in noise levels between biceps and gastrocnemius (p<0.0001, Mann-Whitney test). **b.** *Relationships between mean noise band and optimal AT*_*inc*_. AT_inc_ was calculated from manual counts using 10s windows. The best-fit lines were calculated with weighted least-squares regression due to heteroscedasticity of the data. For biceps (red), *n* was 295 and r^2^ was 0.648. For gastrocnemius (blue), *n* was 304 and r^2^ was 0.402.

#### 2.5.2 Phase two – Use of one-second time windows to refine manual FP counts

By manual inspection, each one-minute recording was categorised as either ‘relaxed’, ‘partially relaxed’, ‘voluntary activity’ or ‘excluded due to poor data quality’. Eighty representative ‘relaxed’ recordings were selected. These underwent a more precise manual FP count using one-second windows. In addition, the time of each FP peak (to the nearest 0.1ms) was recorded. All selected recordings had at least one FP detected manually.

#### 2.5.3 Phase three - Sensitivity, specificity and area under the curve (AUC) of two analytical models

Using the same 80 one-minute recordings from phase two, two analytical models were assessed:

- *Model 1:* this ignored variable noise levels and had a consistent optimal AT_inc_ (Y=A_1_).
- *Model 2:* this used a simple linear model (Y=A_2_X) to describe the relationship between mean noise band and optimal AT_inc_ (*see section 3.2 for justification).*

For each model, the threshold values A_1_ and A_2_ were varied to calculate sensitivities and specificities for each recording. For each threshold, the manual times of each FP peak were compared with the onsets and offsets of all detected FPs and excluded spikes. Automated FPs with a corresponding manual FP were labeled as ‘true positives’. Automated FPs without a corresponding manual FP were labeled as ‘false positives’. Manual FPs without a corresponding automated FP were labeled as ‘false negatives’. Excluded spikes without a corresponding manual FP were labeled as ‘true negatives’. For this validation, other exclusion criteria based on AT_exc_ and IFI were disabled.

A receiver operating characteristic (ROC) curve was produced using the median values of sensitivity and specificity (*figure 3*).(Linden, 2006) An area under the curve (AUC) was calculated for both models as a measure of accuracy. The optimal values for A_1_ and A_2_ were defined as the data-points closest to [0,1]. One hundred repeats of 5-fold cross-validation was performed on pooled data to calculate less biased estimates of the optimal A_2_ value, sensitivity, specificity and classification accuracy.(Kim, 2009, Kohavi, 1995)

**Figure 3.**
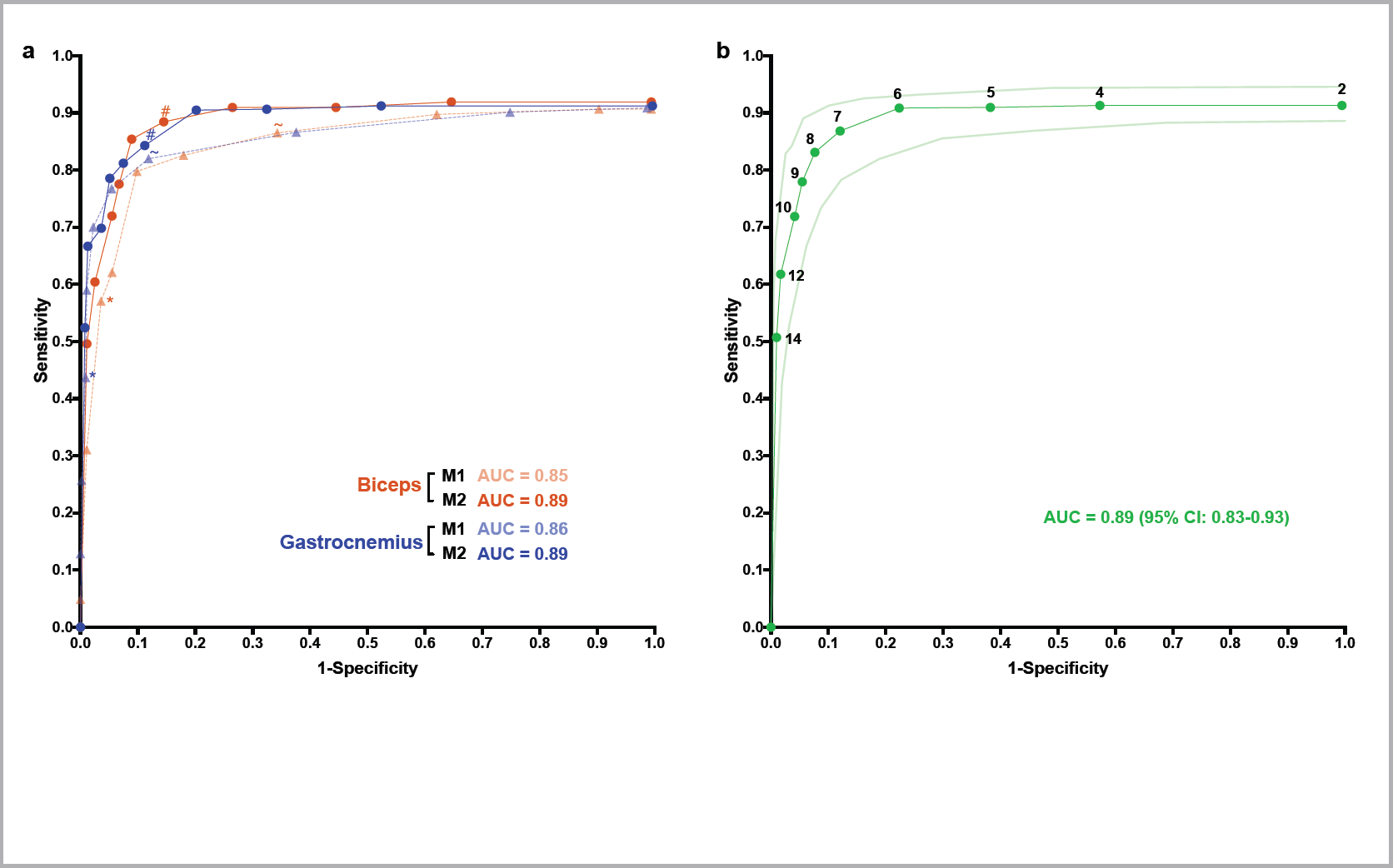
Receiver operating characteristic (ROC) curves for FP identification (phase three). **a** *Comparison of two analytical models.* For each muscle (biceps in red and gastrocnemius in blue), 40 one-minute representative recordings were analysed. Model one (*M1, triangle symbols*) took the form Y=A_1_ and model two (*M2, circular symbols*) took the form Y=A_2_X, where Y was the optimal AT_inc_ (μV), X was the mean noise band (μV) and A_1/2_ was a positive value. From left-to-right, threshold values for A_1_ were 100, 60, 40, 35, 30, 25, 20, 15, 10 and 5, and values for A_2_ were 14, 12, 10, 9, 8, 7, 6, 5, 4 and 2. Median sensitivity and specificity are displayed (non-parametric distributions). Area-under-the curve (AUC, 2 s.f.) represents the accuracy of each model for each muscle. * indicates the performance of M1 when A_1_=40; ~ indicates performance of M1 when A_1_=20; # indicates the performance of M2 when A_2_=7. **b** *Pooled results for model two.* The results from biceps and gastrocnemius were pooled to produce a total of 80 one-minute recordings. Threshold values for A_2_ are displayed. Curves for median sensitivity and specificity are plotted alongside lower and upper limits for 95% confidence interval (CI). AUC represents the accuracy of this model.

#### 2.5.4 Phase four – Amplitude exclusion threshold (AT_exc_)

Each recording from phase three was split into blocks of five seconds duration. A block was excluded if either: a) AT_inc_ exceeded the AT_exc_ for more than half of spikes; or, b) at least one AT_inc_ was greater than double the AT_exc_. If a block were excluded, five seconds were taken from the total time and the FPs detected within it were excluded.

In order to determine the most appropriate AT_exc_, the sensitivity and specificity of 80 recordings were calculated using a range of AT_exc_ (20-100). The lowest AT_exc_ that did not significantly compromise sensitivity was deemed optimal.

#### 2.5.5 Phase five – Comparison of manual and automated FP counts

The optimal model was integrated into the pipeline and used to compare the automated and manual FP counts. A Wilcoxon signed-ranked test was used to detect any significant difference.

## 3 Results

### 3.1 Data processing

The mean number of channels included for further analysis was 19.6 +/-0.4 SEM per recording. The median noise band for all patients was 4.7μV (IQR: 3.2-8.1) for biceps and 2.6μV (IQR: 1.8-4) for gastrocnemius (Mann-Whitney test, p<0.0001; *figure 2*). The one-off pre-processing stage took 10-25 minutes per 30-minute recording. The main analysis took 1-5 minutes per recording for the first run and <20 seconds for subsequent runs.

### 3.2 Optimisation and validation of the analytical pipeline

#### 3.2.1 Phase one - Relationships between manual FP counts, noise levels and optimal amplitude inclusion thresholds

For biceps, a total of 304 one-minute recordings were analysed. Sixteen recordings were excluded due to poor quality. The mean noise bands and corresponding optimal AT_inc_ were plotted (*figure 2b*). Nine outliers were excluded. The best-fit line had slope 11.0 (95% CI 10.1-11.9) and Y-intercept 0.419 (95% CI: −3.01-3.84), forming the equation Y=11.0X.

For gastrocnemius, a total of 312 one-minute recordings were analysed. Eight recordings were excluded due to poor quality. The mean noise bands and corresponding optimal AT_inc_ were plotted (*figure 2b*). Eight outliers were excluded. The best-fit line had slope 17.2 (95% CI 14.9-19.6) and Y-intercept 0.203 (95% CI −5.35-5.75), forming the equation Y=17.2X.

A general relationship of Y=AX existed between mean noise band (X) and optimal AT_inc_ (Y), where A was positive. Although the use of ten-second time windows allowed a large number of recordings to be analysed, we proceeded to phase two, which involved fewer recordings (80) but more precise manual counts over one-second windows.

#### 3.2.2 Phase two – Use of one-second time windows to refine manual FP counts

Out of 320 one-minute biceps recordings inspected manually, 170 (53.1%) were fully relaxed, 42 (13.1%) were partially relaxed, 88 (27.5%) contained only voluntary activity and 20 (6.3%) were of too poor quality to judge. Out of 320 one-minute gastrocnemius recordings inspected manually, 243 (75.9%) were fully relaxed, 46 (14.4%) were partially relaxed, 14 (4.4%) contained only voluntary activity and 17 (5.3%) were of too poor quality to judge.

Eighty representative relaxed recordings were selected, containing 5,318 FPs in total. When counted manually, the median FP frequencies for 40 biceps recordings were 56.5/min (95% CI: 23-84/min) using one-second time windows and 25.5/min (95% CI: 7-47/min) using ten-second time windows. The median FP frequencies for 40 gastrocnemius recordings were 50/min (95% CI: 31-83/min) using one-second time windows and 24.5/min (95% CI: 13-34/min) using ten-second time windows. The differences for each muscle were highly significant (Wilcoxon signed-rank test, p<0.0001).

It was concluded that the time window used to count FPs significantly influenced the manual FP count, which in turn would affect the optimal AT_inc_. Therefore, the specific relationships from phase one were deemed unreliable. Instead, we explored the general relationship (Y=AX) in more detail. We set out to determine the optimal value for A by undertaking ROC analysis.

#### 3.2.3 Phase three - Sensitivity, specificity and AUC of the analytical models

Model 1 ignored variations in noise and took the general form Y=A_1_, where Y was the optimal AT_inc_ (μV) and A_1_ was a positive value. The values of A_1_ that produced the optimal combined sensitivity, specificity and AUC were 30 for biceps and 20 for gastrocnemius (*table 2*). Model 2 accounted for variations in noise and took the general form Y=A_2_X, where Y was the optimal AT_inc_ (μV), X was the mean noise band (μV) and A_2_ was a positive value. The optimal values of A_2_ were 8 for biceps and 7 for gastrocnemius (*table 2*).

**Table 2.** Performance of two analytical models at FP identification. Model 1: Y=A_1_; Model 2: Y=A_2_X, where Y = optimal AT_inc_ (μV), X = mean noise band (μV) and A_2_ = positive value. AUC, area under the curve.

| Model | Muscle | Optimal $A_{1/2}$ | Median sensitivity (%) | Median specificity (%) | AUC (%) |
| --- | --- | --- | --- | --- | --- |
| 1 | Biceps | 30 | 79.8 | 90.2 | 84.1 |
|  | Gastrocnemius | 20 | 82.0 | 89.1 | 84.8 |
| 2 | Biceps | 8 | 85.4 | 91.1 | 88.1 |
|  | Gastrocnemius | 7 | 84.3 | 88.8 | 88.4 |

The performance of model 2 (*figure 3a*) was not significantly different between biceps and gastrocnemius at all thresholds (Mann-Whitney test: all p-values >0.05). This was not the case for model 1, where 8/10 specificities differed between biceps and gastrocnemius (Mann-Whitney test: p-values<0.05). Considering the significant difference in noise levels between the two muscles (Mann-Whitney test: Δ=2.1μV, n=40, p=0.0029), this demonstrated the superior robustness of model 2 when faced with variations in noise levels. This allowed us to pool the data for model 2 (*figure 3b*), on which we performed 100 repeats of 5-fold cross validation. For 49.4% of test folds, the optimal A_2_ value was 8, producing a mean classification accuracy of 87.9% (+/-0.1 SEM), sensitivity of 83.6% (+/-0.2 SEM) and specificity of 91.5% (+/-0.1 SEM). For 50.6% of test folds, the optimal A_2_ value was 7, producing a mean classification accuracy of 85.2% (+/-0.2 SEM), sensitivity of 86.7% (+/-0.2 SEM) and specificity of 85.0% (+/-0.3 SEM). Therefore, due to its superior accuracy, Y=8X was considered the optimal predictive model.

#### 3.2.4 Phase four - Amplitude exclusion threshold (AT_exc_)

Lowering the AT_exc_ beyond 100μV reduced the sensitivity and increased the specificity of the automated model (*figure 4a*). Even when the AT_exc_ was disabled (‘off’), no further increase in sensitivity could be achieved. Therefore, this was a valuable way of excluding excessively noisy portions of data without compromising on the model’s sensitivity.

**Figure 4.**
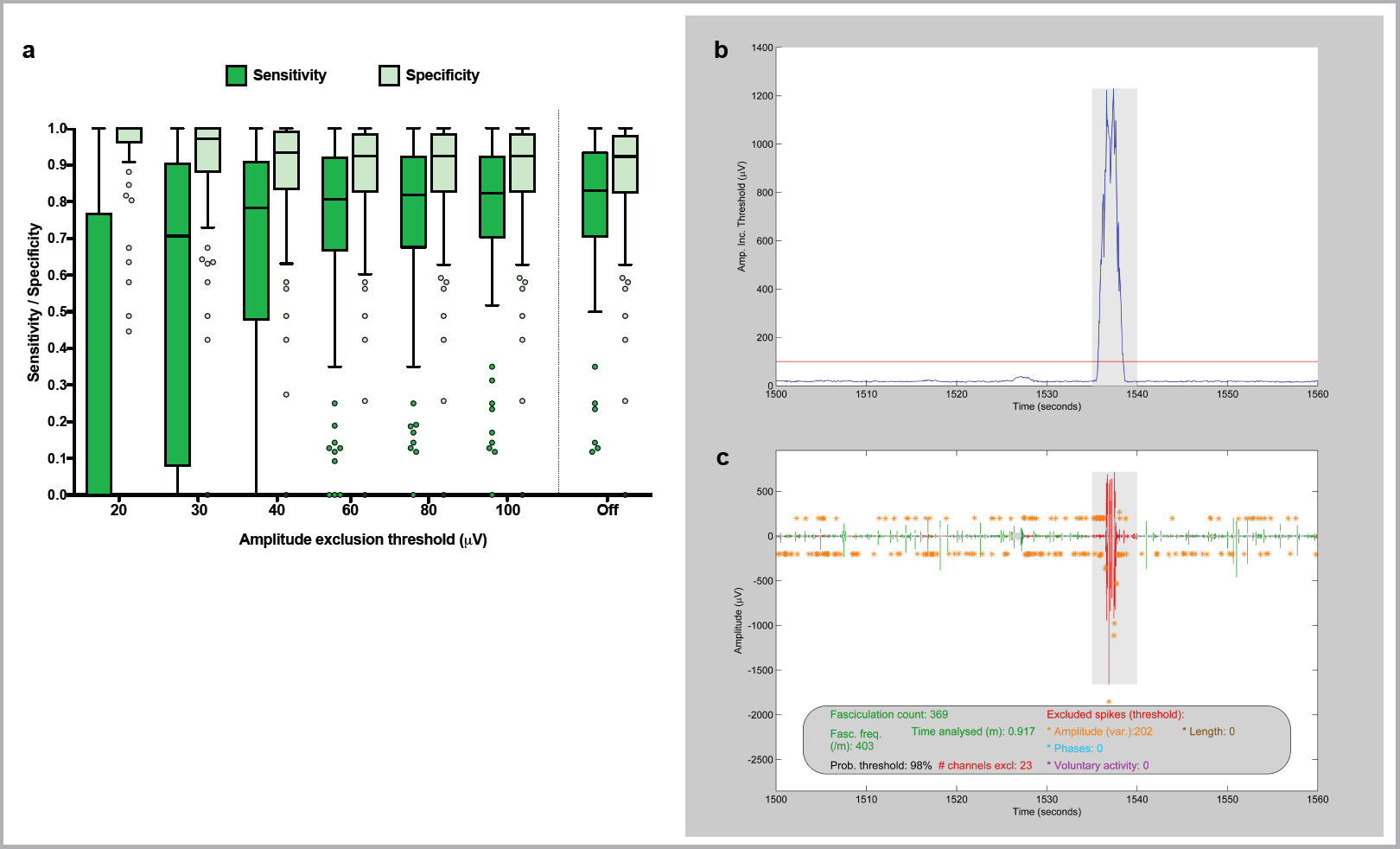
Data exclusion. **a** *Finding the optimal amplitude exclusion threshold (AT*_*exc*_). The *AT*_*exc*_ was varied to calculate the sensitivity and specificity for 80 pooled (biceps and gastrocnemius) one-minute recordings. Boxes represent median and inter-quartile range (IQR). Whiskers represent (upper quartile + 1.5*IQR) and (lower quartile −1.5*IQR) according to Tukey’s method. Data points beyond this range are plotted individually. **b** *AT*_*exc*_ *example - part 1.* One minute of amplitude inclusion thresholds (linearly related to noise levels), showing a burst in noise levels between 1535-1540s. Amplitude exclusion threshold of 100μV (red line) applied to exclude shaded region. **c** *AT*_*exc*_ *example - part 2.* Corresponding one minute of super-channel recording, showing exclusion of noisy period. Fasciculation count (369), time analysed (0.917m) and fasciculation frequency (403/m) have been automatically adjusted.

#### 3.2.5 Phase five – Performance of the optimal model

The optimal automated method was model 2 (Y=A_2_X), where A_2_ = 8 and AT_exc_ = 100 (*figure 5*). For 80 recordings with high precision FP counts, there was no significant difference between the manual and automated counts (Wilcoxon signed-rank test, p=0.394).

**Figure 5.**
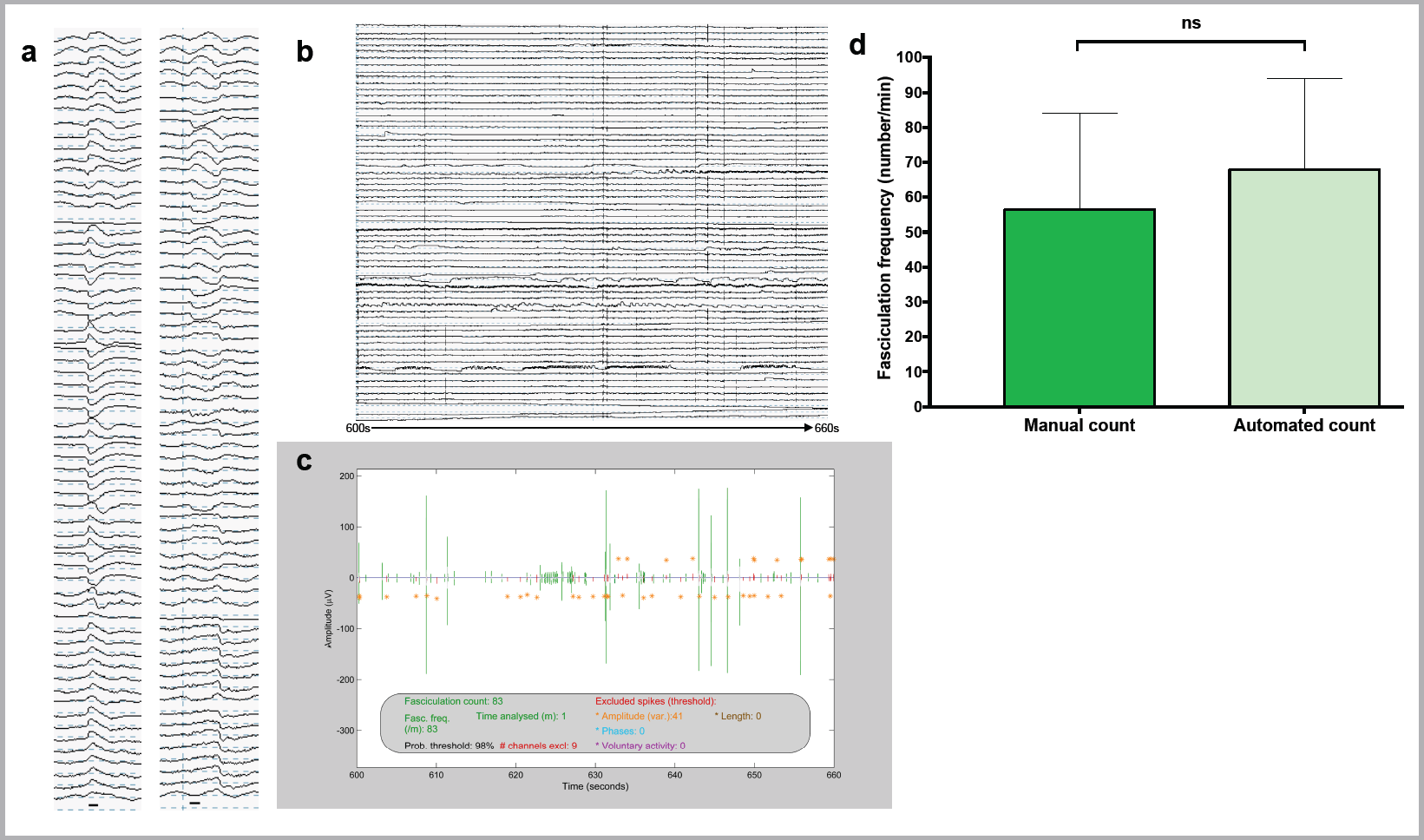
Application of the analytical pipeline. **a** *FPs from raw data.* Black bar indicates 5ms duration. **b** *Example raw data.* 60s of raw data from gastrocnemius of ALS patient 3. **c** *Example SPiQE output.* Super-channel of corresponding 60s of data in ‘b’ after application of optimal analytical pipeline. **d** *Model performance.* The optimal model (Y=8X) has been applied to 80 one-minute recordings and compared with high precision manual counts. Medians and 95% confidence intervals plotted. Wilcoxon signed-rank test confirms no significant difference (p=0.394).

## 4 Discussion

SEMG has improved the detection rate of fasciculations compared to clinical examination or NEMG.(Howard and Murray, 1992, Hjorth et al., 1973) The advent of HDSEMG, where multiple channels are aligned in a linear or grid formation, has provided superior spatial resolution and muscle coverage compared to single channel SEMG.(van Dijk et al., 2010) SPiQE combines this non-invasive sensor technology with an innovative signal-processing algorithm to quantify FPs in an accurate and automated way. Although other groups have applied automated techniques to FP analysis,(Drost et al., 2007, Jahanmiri-Nezhad et al., 2014b) we aimed to combine relatively simple signal-processing methods to convert unrefined HDSEMG recordings into reliably interpretable results.

The main challenge facing this automated analytical model concerned fluctuating noise levels.(De Luca et al., 2010) Biceps produced higher noise levels than gastrocnemius, prompting us to validate these muscles separately. The fact that the optimal models from each muscle were not significantly different from each other demonstrated the robustness of the analytical technique, which allowed us to pool the data to develop a unified model.

An established method for calculating noise in EMG data is the root mean square.(Hug et al., 2006, Jahanmiri-Nezhad et al., 2014a) However, we did not feel this was the optimal approach in this setting. We defined the noise band to be directly comparable with the peak-trough amplitudes of potential fasciculations. A single noise band represented the average peak-trough noise within the immediate vicinity of its corresponding spike and could be calculated without significant computational expense.

Importantly, the empirical relationship between noise band and optimal amplitude inclusion threshold was linear and the Y-intercept did not significantly differ from zero, leading to a simplification of the enhanced model. This latter observation seems intuitive when considering the hypothetical scenario of zero noise, in which case any amplitude deviation from zero would be a true signal. Not only was model 2 the most accurate, but it was also the most robust to shifts in background noise (*figure 4a*). Like others, our original approach had been to apply the same amplitude inclusion threshold across every recording.(de Carvalho and Swash, 2013, de Carvalho and Swash, 2016a) Anecdotally, we observed major problems with this approach, particularly in recordings with higher noise levels. By testing models 1 and 2 in parallel on the same representative recordings, we were able to confirm the superiority of a noise-responsive model.

An interesting feature of the validation, as evidenced in *figure 4*, was that the sensitivity reached a saturation point with reducing A_2_ values. Our interpretation of this relied on understanding that false negatives in this context could be divided into two groups. In the first group were those events detected as spikes but whose amplitudes fell short of the amplitude inclusion threshold. The second group represented those that were not recognised as spikes in the first place, due to a probability threshold (set at 98%) that was not sufficiently inclusive. The observed saturation level in sensitivity was produced by the second group. This indicated a key avenue to explore to improve the model’s accuracy in future iterations.

We estimated the accuracy of the two models using ROC analysis, adopted from its widely used role in clinical diagnostics.(Linden, 2006) To establish the ground truth, we developed a simple definition of a fasciculation potential for manual analysis, providing a standardised approach across all recordings. More than one assessor for manual counts would be beneficial,(Harding et al., 2016) however this is very time-consuming. The single assessor (JB) had accumulated the most experience in our group analysing raw HDSEMG data manually.

Our analytical models have been trialed in a relatively small group of patients, however the inclusion of multiple visits, multiple muscles and control patients adds to the robustness of the optimal model. An internal cross-validation approach was used to provide less biased predictive parameters, however application in an external group of patients would enhance its validity.

We have focused on biceps and gastrocnemius as their size permits the use of large HDSEMG grids, thereby maximising data coverage for the purposes of validation and interpretation. Our decision to study gastrocnemius, a muscle known to produce prominent fasciculations in the healthy population,(Fermont et al., 2010, Simon and Kiernan, 2013) reinforces our aim to identify pathophysiological features that discriminate disease and healthy states. Moreover, these two muscles were the focus of a complementary automated analytical tool using ultrasound.(Harding et al., 2016) Ultrasound can provide diagnostic and practical advantages over EMG, suggesting the non-invasive combination of ultrasound and HDSEMG warrants exploration as we strive to understand the electromechanical properties of fasciculations.(Johansson et al., 2017, Tsuji et al., 2018)

Fasciculations are a hallmark clinical feature in ALS and reflect early neuronal dysfunction so could provide a sensitive measure of motor neuron health. SPiQE was conceived as a means to understand the natural history of fasciculations in relation to disease progression, anticipating that this could be translated into a novel outcome measure in therapeutic trials. This validation is a promising introduction, paving the way for larger prospective studies. Although others have not found evidence that FP frequency correlates with neurological decline in ALS,(de Carvalho and Swash, 2016a) we felt that a more comprehensive approach based on an automated analytical method may be more fruitful. The aim would be to prospectively compare FP parameters, such as frequency or amplitude, with established markers of disease progression, including the ALS-Functional Rating Scale (ALS-FRS) and Motor Unit Number Index (MUNIX).(Cedarbaum et al., 1999, Neuwirth et al., 2015, Neuwirth et al., 2017)

## 5 Conclusion

SPiQE is the first attempt at an automated tool to comprehensively identify fasciculations in ALS and related disorders using unrefined HDSEMG. Performing robustly amidst fluctuating noise levels, it achieves a favourable average classification accuracy of 88%. Instead of relying on complex theoretical concepts, its design has been guided by real-world data collected prospectively from patients and controls. It is anticipated that this empirical approach will improve its translation into a meaningful clinical tool. Our multi-disciplinary group has combined expertise from bioengineering, clinical neurology and biostatistics - a combination that’s led to the customisation and simplification of the technical approach. Throughout our clinical study program, SPiQE will serve as a platform to discover potential correlations between pathological FP parameters and neurological decline in patients with ALS, thereby aiding the search for a novel disease biomarker.

## Abbreviations

ALS, amyotrophic lateral sclerosis; AT_exc_, amplitude exclusion threshold; AT_inc_, amplitude inclusion threshold; AUC, area under the curve; BFS, benign fasciculation syndrome; FP, fasciculation potential; (HD)SEMG, (High-density) surface electromyography; IFI, inter-FP interval; MMN, multifocal motor neuropathy; MU, motor unit; NEMG, needle electromyography; ROC, receiver operating characteristic; SEM, standard error of the mean; SPiQE, Surface Potential Quantification Engine.

## Author contributions

JB setup the study and collected the data. JB and AW devised and tested the analytical technique. EM, MB, KM and CS supervised the project and provided expert technical and clinical guidance. RI provided expert input on statistical learning. JB wrote the article with editing from co-authors.

## Acknowledgements

J. Bashford was supported for 14 months by the Sattaripour Charitable Foundation and the Motor Neurone Disease Association (Shaw/Jul15/932-794). Ongoing funding for 36 months is provided through the MRC/MNDA Lady Edith Wolfson Clinical Research Training Fellowship (MR/P000983/1). A. Wickham was funded by a PhD studentship in the EPSRC Centre for Doctoral Training in Neurotechnology for Life and Health. We would like to thank all the patients involved in this study for their willingness and determination to participate. We thank TMSi for supplying the amplifier and sensors.

